# Integrative Computational Analysis of VCX2 in Hepatocellular Carcinoma: From Potential Biomarker Discovery to Therapeutic Targeting with Peruvian Natural Products

**DOI:** 10.1101/2025.02.20.639333

**Authors:** Luis Daniel Goyzueta-Mamani, Haruna Luz Barazorda-Ccahuana, Mayron Antonio Candia-Puma, Nadia M. Hamdy, Miguel Angel Chávez-Fumagalli

**Affiliations:** Computational Biology and Chemistry Research Group, Vicerrectorado de Investigación, Universidad Católica de Santa María, Arequipa 04000, Peru; Facultad de Ciencias Farmacéuticas, Bioquímicas y Biotecnológicas, Universidad Católica de Santa María, Arequipa 04000, Peru; Biochemistry Department, Faculty of Pharmacy, Ain Shams University, Abassia, Cairo 11566, Egypt

**Keywords:** Hepatocellular carcinoma (HCC), single-cell RNA sequencing, VCX2, cancer biomarker, Luteolin-5-O-glucoside, Computational Analysis, Natural products

## Abstract

Hepatocellular carcinoma (HCC) is a leading cause of cancer-related mortality, often developing in the context of chronic liver disease, fibrosis, and cirrhosis. Identifying novel biomarkers with diagnostic and therapeutic potential is essential, particularly those relevant across multiple cancer types. This study integrates single-cell RNA sequencing (scRNA-seq) data from healthy and diseased liver tissues, analyzing different cellular lineages to identify genes involved in fibrosis, angiogenesis, immune modulation, and apoptosis regulation. Uniform Manifold Approximation and Projection (UMAP) clustering, differential gene expression (DEG) analysis, and protein-protein interaction (PPI) network construction were employed to identify genes contributing to tumor progression and metabolic reprogramming. Key genes, including Transmembrane BAX Inhibitor Motif Containing 4 (TMBIM4), Regulator of G-protein signaling 5 (RGS5), CEA Cell Adhesion Molecule 7 (CEACAM7), and Variable Charge X-Linked 2 (VCX2), exhibited significant roles in tumorigenesis and chromosomal stability. VCX2, a cancer/testis antigen, emerged as a potential biomarker and druggable target due to its altered expression among multiple cancers. Structural modeling and molecular docking (MD) of VCX2 identified a high affinity binding pocket, guiding a virtual screening of Peruvian natural products. Luteolin-5-O-glucoside, from *Equisetum arvense*, was identified as the most promising compound, showing a strong docking score (−7.42 kcal/mol) and favorable binding free energy (ΔG_bind = −40.13 kcal/mol). MMGBSA calculations revealed stabilizing hydrogen bonds with PRO91, GLU97, and GLU109, reinforcing its strong binding stability. These findings position VCX2 as a promising target for HCC therapy and suggest Luteolin-5-O-glucoside as a lead compound with high drug-like potential. Further studies should focus on experimental validation, molecular dynamics simulations, and structure-activity relationship (SAR) optimization to advance VCX2-targeted therapies.

**Highlight statements:**

- VCX2 as a Biomarker exhibited differential expressions in HCC versus healthy liver tissue and a suggested role in tumor progression and chromosomal stability.
- Luteolin-5-O-glucoside from *Equisetum arvense* was identified as a promising compound: strong docking score (−7.42 kcal/mol), favorable binding free energy (ΔG_bind = −40.13 kcal/mol), and stabilized interactions with key amino acids (PRO91, GLU97, GLU109).
- VCX2 may serve as an oncogenic driver; small molecule inhibition could desensitize tumor cells that need further refinement and validation of structural models due to lack of experimentally resolved crystal structure.
- VCX2 is a novel biomarker and drug target for HCC with Luteolin-5-O-glucoside presents potential for targeted therapy, paving the way for precision medicine approaches.

## 1. Introduction

Hepatocellular carcinoma (HCC) is the predominant primary malignancy of the liver, representing approximately 75-85% of all liver cancer cases globally, with approximately 830,000 fatalities annually [1,2]. It is a significant public health challenge due to its rising incidence, high mortality rate, and complex etiology. HCC typically develops in association with chronic liver disease, with hepatitis B virus (HBV) and hepatitis C virus (HCV) infections being the leading risk factors worldwide [3,4]. Other etiological factors comprise alcohol abuse, non-alcoholic fatty liver disease (NAFLD), aflatoxin exposure, and metabolic disorders. These risk factors lead to chronic inflammation, fibrosis, and eventual cirrhosis, which is observed in 80-90% of HCC cases at the time of diagnosis [5].

The global burden of HCC is significant, with an estimated 905,000 new cases and 830,000 deaths annually, making it the sixth most common cancer and the third leading cause of cancer-related deaths worldwide [6]. The geographic variation in HCC incidence reflects differences in risk factor prevalence. Regions such as East Asia and Sub-Saharan Africa, characterized by high HBV endemicity, exhibit the highest incidence rates [7]. In contrast, Western countries have experienced an increase in cases related to the growing prevalence of NAFLD and obesity [8].

The molecular development and progression of cancer involve diverse genetic and epigenetic alterations. Novel biomarkers, including VCX2, TMBIM4, RGS5, and Tropomyosin Beta Chain 4X (TM5B4X), can potentially advance our understanding of tumor biology. VCX2 is an X-linked gene predominantly expressed in testicular germ cells and classified as a cancer/testis antigen (CTA) [9]. Typically silenced in normal somatic tissues, it can be aberrantly reactivated in certain malignancies, where it may contribute to tumor progression through immune evasion and altered gene regulation. TMBIM4, a member of the BCL2 associated X, apoptosis regulator (BAX) inhibitor family, is crucial for apoptosis regulation, acting as a negative regulator of programmed cell death. It enhances cancer cell survival by modulating mitochondrial function and calcium homeostasis, leading to resistance against apoptosis-inducing therapies [10]. RGS5 is involved in G-protein signaling and plays a key role in angiogenesis and vascular remodeling. Its upregulation in tumors promotes the formation of abnormal vasculature, which supports tumor growth and immune escape, making it a potential target for anti-angiogenic therapies [11]. TM5B4X, part of the tropomyosin family, is critical for cytoskeletal organization and cell motility. Dysregulation of TM5B4X has been linked to increased metastatic potential in various cancers, as it facilitates actin filament remodeling, aiding in cancer cell migration and invasion [12]. These biomarkers highlight the complexity of tumor heterogeneity and offer opportunities for targeted therapeutic strategies. In recent years, the utilization of natural products has gained interest as a complementary strategy to combat HCC by modulating cancer-specific biomarkers and pathways. Natural compounds such as curcumin, resveratrol, quercetin, prodigiosin, and berberine have demonstrated anti-cancer effects, including inhibition of angiogenesis, suppression of cell survival pathways, and induction of apoptosis [13,14]. Curcumin can inhibit TMBIM4-mediated resistance to apoptosis [15], while resveratrol efficiently targets angiogenesis-related pathways, potentially impacting RGS5 activity [16]. These natural agents demonstrate multi-targeted effects and hold the potential to mitigate the adverse effects of conventional therapy.

The therapeutic application of natural products originates from ancient medical traditions, wherein plant-derived compounds were central to treating liver-related disorders. Traditional Chinese Medicine (TCM) and Ayurveda have historically utilized herbs including *Phyllanthus niruri*, licorice root, and milk thistle for their hepatoprotective properties [17]. Modern pharmacological research has validated many of these applications, demonstrating their capacity to influence molecular pathways associated with HCC. For instance, milk thistle’s active component, silymarin, has been shown to inhibit tumor growth and proliferation by modulating oxidative stress and inflammatory signaling [18].

Clinically, HCC is frequently asymptomatic in its early stages, leading to late-stage diagnoses that limit treatment alternatives [19]. Surveillance programs utilizing imaging techniques and conventional biomarkers, such as alpha-fetoprotein (AFP), have been established for high-risk populations [20]. However, incorporating natural compounds that target specific biomarkers like VCX2, TMBIM4, RGS5, and TM5B4X offers an innovative approach to enhancing early intervention and therapy. This study aims to identify a potential biomarker in HCC, by integrating scRNA-seq data from healthy and diseased liver tissues, and differential gene expression analysis and protein-protein interaction networks. Structural modeling, MD, and virtual screening using the Peruvian Natural Products Database (PeruNPDB) assess VCX2’s druggability and potential inhibitors. This study provides a computational framework for biomarker discovery, emphasizing the need for experimental validation and structural characterization.

## 2. Material and Methods

### Single-cell RNA sequencing data collection and processing

The gene raw counts or normalized gene expression matrix of single-cell gene expression data from the liver of 5 relatively healthy patients (Accession No. GSE115469) [21], were downloaded from the Gene Expression Omnibus (GEO) (https://www.ncbi.nlm.nih.gov/geo/) [22]. The cancer tissues data was searched at the CancerSCEM database (https://ngdc.cncb.ac.cn/cancerscem/index) [23], where HCC from 2 samples [24]; Muscle-invasive Urothelial Bladder Cancer from 1 sample [25]; Triple Negative Breast Cancer from 6 samples [26]; Breast Ductal Carcinoma in situ from 1 sample [26]; Colorectal Cancer from 16 samples [27]; Lung Adenocarcinoma from 21 samples [28]; Lung Squamous Cell Carcinoma from 7 samples [29]; Non-small Cell Lung Cancer from 7 samples [29]; Ovarian Carcinoma from 4 samples [30]; Pancreatic Ductal Adenocarcinoma from 24 samples [31] and Stomach Adenocarcinoma from 1 sample [32] were selected to constitute the organs dataset.

### Data Analysis

The data was loaded into R 4.1.0 and was preprocessed using standard parameters of the R packages ‘Seurat’ v.4.0.3 [33]. Briefly, the expression matrix for each dataset was merged into one Seurat object using the *“CreateSeuratObject”* function. At the same time, cells with less than 100 expressed genes and higher than 25% mitochondrial genome transcript and genes expressed in less than three cells, were removed. The gene expression data was normalized using the *“NormalizeData”* function. The sources of cell-cell variation driven by batch were reverted, using the number of detected UMI and mitochondrial gene expression by the *‘‘ScaleData’’* function. The *“FindVariableGenes”* function identified highly variable genes and was used for the principal component analysis (PCA) on the highly variant genes using the *‘‘RunPCA’’* function. The “JackStraw” function was implemented to remove the signal-to-noise ratio. Cells were then clustered utilizing the *‘‘FindClusters’’* function by embedding cells into a graph structure in PCA space. The clustered cells were then projected onto a two-dimensional space using the *“RunUMAP”* function. To create merged datasets of different organs, the *“MergeSeurat”* function was applied; the raw count matrices of two Seurat objects or more were merged into one; and a new Seurat object was created with the resulting combined raw count matrix.

Differentially expressed genes (DEGs) were identified by the *“FindAllMarkers”* or *“FindMarkers”* function, applying the Wilcox test, which returned “p_val_adj” using the Bonferroni correction and the log-transformed fold change, “avg_logFC”. The resulting cell clusters were annotated for cell type identification using the *“SingleR”* package, which also computes the expression correlation scores between each test cell and each cell type in the reference cell atlas. The reference cell type with the highest correlation serves as the basis for cell identity [34]. Gene Ontology (GO), regarding Biological Processes (BP), Molecular Functions (MF) and Cellular Components (CC), and Kyoto Encyclopedia of Genes and Genomes (KEGG) pathway and enrichment analyses were performed with *“clusterProfiler”* [35] and *“enrichplot”* [36] R packages. UMAP dimensional reduction plots, bar plots, volcano plots, dot plots, and expression and enrichment heatmaps were generated with the *“SCpubr”* package [37].

### Protein-Protein Interaction (PPI) Network Analysis

The selected sequences were chosen for the PPI network building. The molecular networks were retrieved from the Search Tool for the Retrieval of Interacting Genes/Proteins (STRING) database version: 12.0 (https://string-db.org) [38] and analyzed at the Cytoscape platform [39], where the plugin cytoHubba was used to score and rank the nodes according to network properties, affording to the Maximal Clique Centrality (MCC) topological analysis method [40]. Also, the plugin StringApp [41] was used to retrieve functional enrichment for GO terms, BP, MF, and CC, and for accessing the TISSUES database v2.0 of human tissue expression data [42]. Cytoscape default settings were considered to visualize the network, whereas node size and color were manually adjusted, considering the scores provided by the analysis.

### Protein Preparation

To refine the VCX2 structural model, which exhibited misconformations in the AlphaFold prediction, molecular dynamics simulations were first performed to correct structural deviations and stabilize the protein before docking studies. This was followed by structural optimization using Schrödinger Maestro to refine the druggable binding pocket along specific X, Y, and Z coordinates.

The molecular dynamics simulations followed a four-step protocol: energy minimization, annealing, equilibration, and production simulation. First, energy minimization was conducted using 5,000,000 iterations of the steepest descent integrator to eliminate steric clashes. Next, simulated annealing was applied over 12 cycles (500 ps each), gradually heating the system to 500 K and cooling it to 300 K over 50 ns, enhancing conformational sampling and correcting structural misfolding. Equilibration was performed under the NVT ensemble at 300 K for 5 ns, applying position restraints on the protein backbone while allowing the solvent to adapt. Finally, a production simulation under the NPT ensemble was executed at 309 K and 1 bar for 1000 ns, ensuring a stable and reliable VCX2 structure suitable for downstream docking analyses. After MD stabilization, the refined structure underwent further optimization in Schrödinger Maestro (v.12.8.117). Using the Protein Preparation Wizard, bond orders were assigned (CCD database), missing hydrogen atoms were added, and loops/side chains were reconstructed (Prime module). The hetero state at pH 7.0 was generated using Epik, and disulfide bonds were preserved at zero bond orders. All water molecules, solvents, and ligands were removed to prepare a clean docking environment. Hydrogen bonds were optimized, and energy minimization was applied using the OPLS4 force field, excluding water molecules beyond 3.0 Å from heteroatoms. This final step refined the druggable pocket along specific X=31.42, Y=50.12, and Z=41.28 coordinates (size 10Å), ensuring structural accuracy for MD and ligand interaction studies.

### Virtual screening with the Peruvian Natural Products Database (PeruNPDB)

The simplified molecular-input line-entry system (SMILES) [43] of Peruvian natural products previously described were retrieved from the PeruNPDB online web server (https://perunpdb.com.pe/), which contains information on 280 molecules from Peruvian biodiversity [44]. Furthermore, the compounds were imported into OpenBabel within the Python Prescription Virtual Screening Tool [45] and subjected to energy minimization. PyRx performs structure-based virtual screening by applying docking simulations using the AutoDock Vina tool [46]. The targets were uploaded as macromolecules, and a thorough search was carried out by enabling the “Run AutoGrid” option, which creates configuration files for the grid parameter’s lowest energy pose, and then the “Run AutoDock” option, which uses the Lamarckian GA docking algorithm. The docking simulation was then run with an exhaustiveness setting of 20 and instructed only to produce the lowest energy pose. The Z-score was calculated for the dataset and the results were uploaded within the GraphPad Prism software version 10.0.2 (232) for Windows from GraphPad Software, San Diego, California, USA, at http://www.graphpad.com, and plotted as a violin plot, while a critical Z-score value of ±1.645 for a significance level of 0.05, was selected as cut off value for positive binding.

### Ligand Preparation, Active Site Calculation, and Glide Grid Generation

The best ligand was prepared using the LigPrep wizard, applying Epik at pH 7.0 (±2) to generate probable protonation states and tautomers, while maintaining a maximum of 32 stereoisomers per ligand to ensure comprehensive conformational sampling. Following protein optimization, the active sites on the 3D models were identified using the SiteMap module in the Schrödinger suite. This process involved detecting top-ranked potential receptor binding sites, with site maps cropped at 4 Å from the nearest site point, resulting in five predicted active sites. For docking studies, the Receptor Grid Generation wizard was used to create a grid file centered on the highest-scoring active site. The grid box size was adjusted to ensure full coverage of the binding pocket, optimizing the docking process. This refined grid was then used for molecular modeling simulations to evaluate ligand interactions within the active site.

### Molecular Docking (MD)

With the active site refined, docking simulations were performed using the Glide module in Schrödinger, employing a three-tiered docking approach: High-Throughput Virtual Screening (HTVS) for rapid ligand filtering, Standard Precision (SP) for balanced accuracy and efficiency, and Extra Precision (XP) for detailed evaluation of top-ranked ligands. The docking scores were analyzed to assess binding affinity and interaction strength, with ligand-protein interactions visualized using Maestro’s interaction diagram tool before proceeding to molecular modelling simulations for further validation.

### Prime MM-GBSA

The Prime MMGBSA method (Prime Version 4.8) was used to calculate the relative binding free energy (ΔG_bind) of the ligand molecule. The formula used for this calculation is provided below:

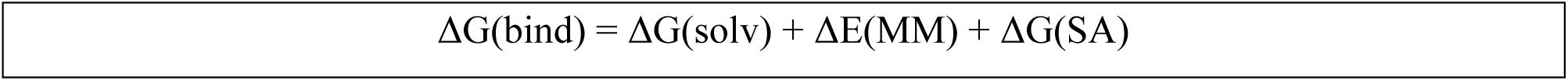

where: ΔGsolv is the difference in GBSA solvation energy of the protein-ligand complex and the sum of the solvation energies for unliganded protein and ligand. ΔEMM is a difference in the minimized energies between the protein-ligand complex and the sum of the energies of the unliganded protein and ligand. ΔGSA is a difference in the surface area energies of the complex and the sum of the surface area energies for the unliganded protein and ligand.

Prime MM-GBSA computes the energy of the optimized free receptor, free ligand, and the ligand-receptor complex. Additionally, it determines the ligand strain energy by simulating the ligand in a solvated environment generated using the VSGB 2.0 suite. The Prime Energy Visualizer was used to visualize energy distributions and interactions.

## 3. Results and Discussions

HCC is among the top causes of cancer-related mortality worldwide, frequently developing in the setting of chronic liver disease, fibrosis, and cirrhosis [47]. The search for novel cancer biomarkers is crucial for improving diagnostic precision and therapeutic strategies [48]. This study utilizes available resources to analyze gene expression in healthy and diseased liver tissues, focusing on genes expressed in cell lines associated with liver pathology and cancer. By integrating scRNA-seq data from healthy liver tissue with datasets from various cancers, we aim to identify genes that function as liver-specific disease markers, thereby clarifying their roles in tumor progression and potential applications in precision medicine.

Our research focused on HCC and its wider application to other cancers to find potential biomarkers that could differentiate between healthy and diseased liver states. Specifically, we aimed at the expression of genes implicated in processes like fibrosis, angiogenesis, immunological modulation, and apoptosis regulation within cellular lines. Our analysis began with scRNA-seq data, which offered a comprehensive picture of transcriptional heterogeneity across liver cell types and served as the starting point for the research.

The UMAP plot in *Figure 1A* shows distinct clusters with different transcriptional fingerprints. Each cluster is identified by color and represents a certain cell type or functional state. The clustering highlights the diversity of liver cell types, with each cluster representing a specific lineage or biological function. A more detailed breakdown of the color representation is provided in *Figure 2A*. The overlay of healthy (green) and diseased (red) liver samples in *Figure 1B* illustrates significant transcriptional shifts and population changes during disease progression. Diseased liver samples show a higher prevalence of certain clusters, reflecting fibrosis, immune infiltration, and metabolic dysregulation associated with disease progression.

**Figure 1.**
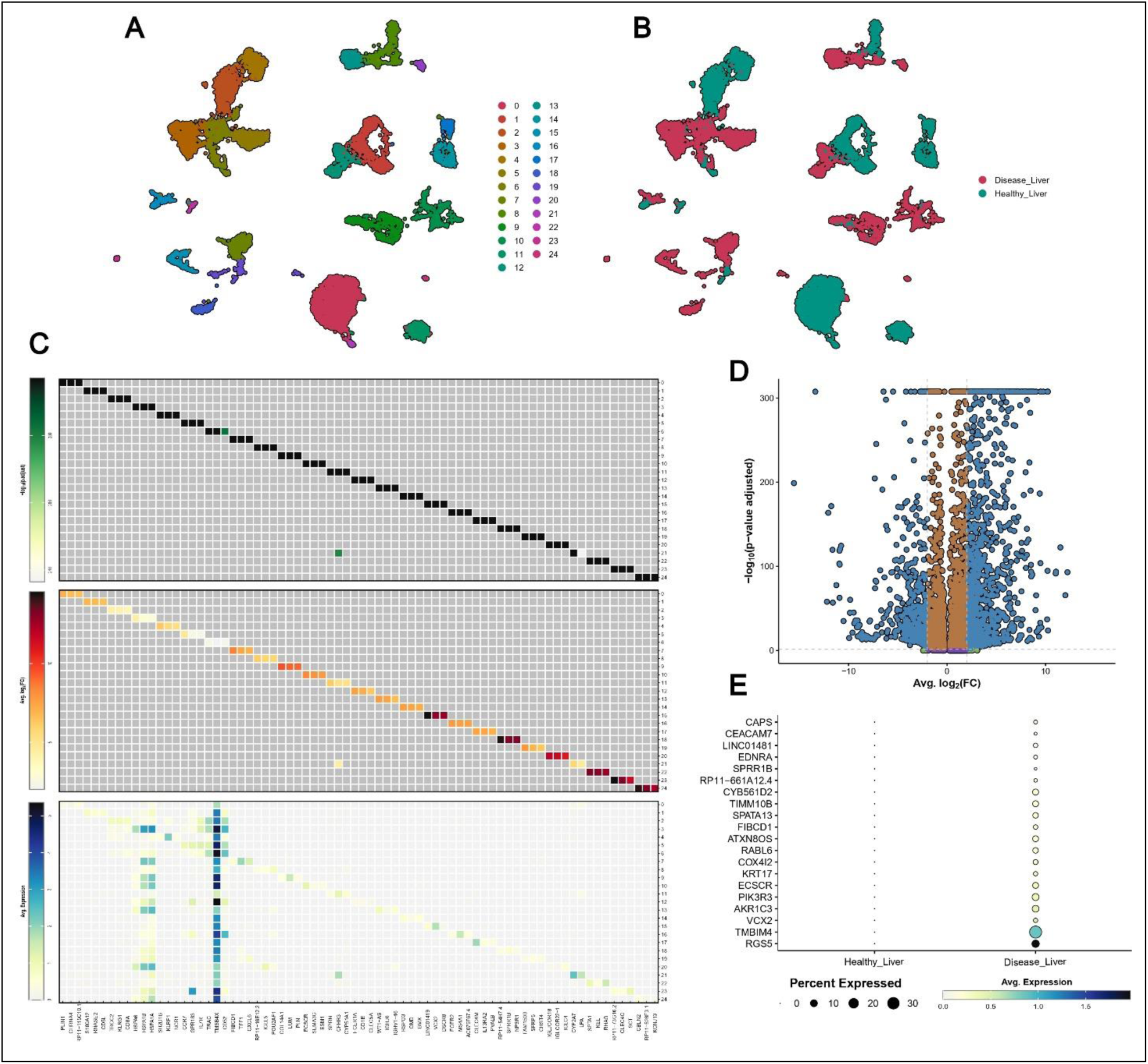
Comprehensive analysis of liver cell populations and transcriptional changes in healthy and diseased liver tissues. (A) UMAP plot displays distinct liver cell clusters, color-coded by cell type. Healthy liver is dominated by hepatocytes (green), while diseased liver exhibits a marked expansion of fibroblasts (brown), macrophages (blue), and immune cells, including T cells (dark pink) and B cells (light pink), reflecting cellular reorganization during disease progression. (B) UMAP overlay of healthy (green) and diseased (red) liver samples highlights the transcriptional shifts and changes in cellular composition associated with liver cancer. (C) Heatmaps illustrate gene relationships and expression changes, with the upper heatmap showing gene correlations among the top 3 markers per cluster. (D) Volcano plot depicts differentially expressed genes between healthy liver and HCC, highlighting upregulated genes critical to cancer progression. (E) The dot plot visualizes average expression (color intensity) and percent of cells expressing key genes (dot size) across healthy and diseased liver cell types.

**Figure 2.**
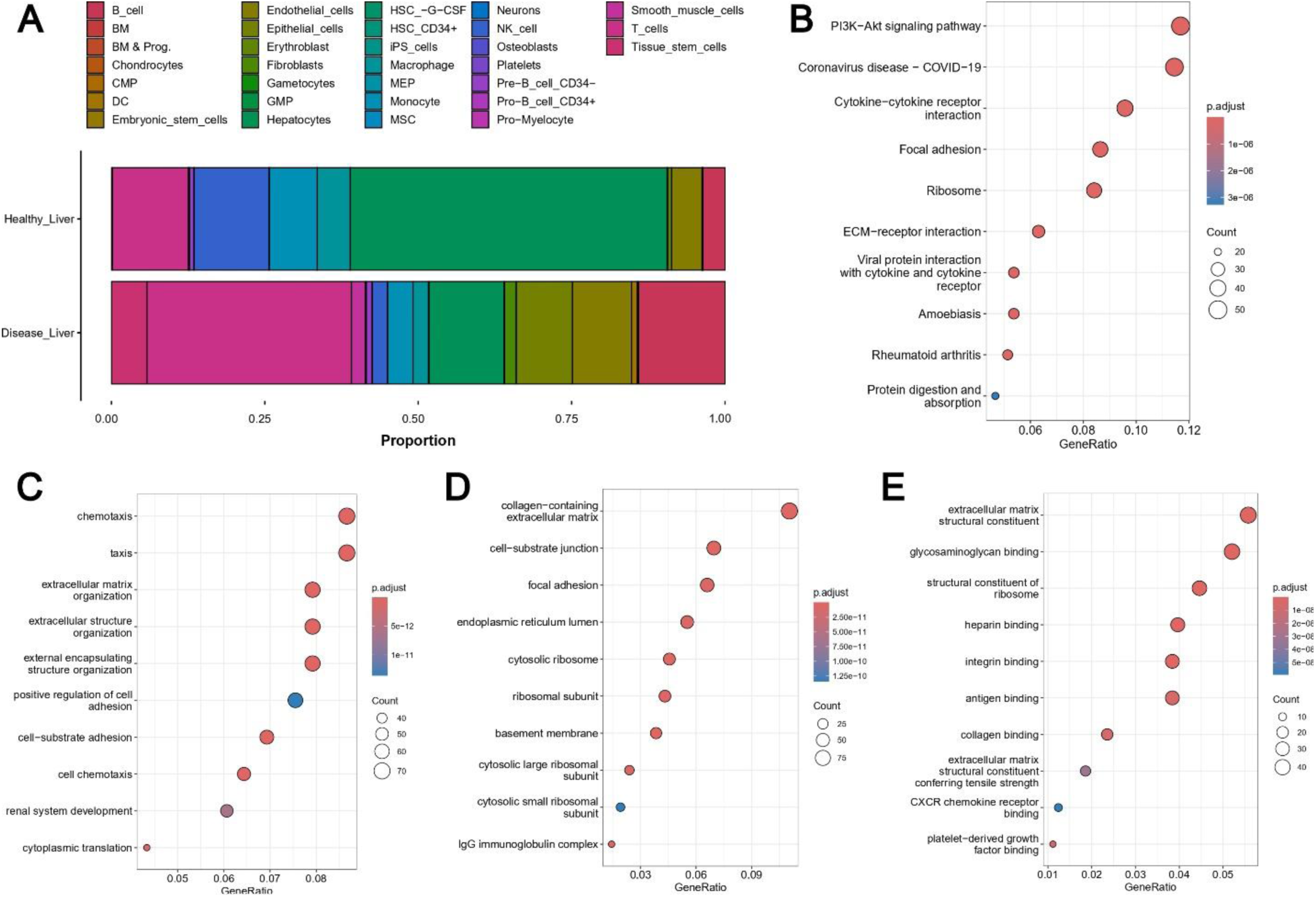
Comprehensive analysis of molecular, cellular, and functional alterations in healthy and diseased liver tissues. (A) Proportional representation of cell types in healthy and diseased liver tissues. (B) KEGG pathway enrichment analysis. (C) Biological process enrichment. (D) Cellular component enrichment. (E) Molecular function enrichment.

Figure 1C presents a heatmap illustrating the relationships among the top three markers per cluster, focusing on the expression patterns. This heatmap consists of three sections: the upper heatmap represents the correlation matrix, the central heatmap depicts statistical significance (p-values), and the lower heatmap highlights the log2 fold-change in gene expression. Together, these components offer a comprehensive view of marker expression differences between healthy and diseased liver tissues. The correlation matrix (upper heatmap) reveals genes co-expression within the clusters and shared regulatory pathways. For instance, TMBIM4 and RGS5 demonstrate a strong correlation, reflecting their roles in apoptosis inhibition and vascular remodeling [10]. These markers are closely linked to cancer progression due to their contributions to tumor survival, angiogenesis, and stress adaptation [49]. These interactions suggest that these genes may co-regulate important pathways in HCC. The central heatmap illustrates p-values and highlights statistically significant differences in gene expression between healthy and diseased liver tissues. Significant markers include TMBIM4, RGS5, and CEACAM7, which are particularly interesting due to their particular expression profiles and function in liver cancer. For instance, CEACAM7, a cell adhesion molecule, is known for its tumor-suppressive properties in healthy tissues but can exhibit altered expression during tumorigenesis, indicating its dual role in liver pathophysiology [50].

The lower heatmap, representing log2 fold-change, provides quantitative insights into the extent of gene expression changes. Genes like TMBIM4, RGS5, and Cytochrome B5 Reductase 3 (CYB5R3) are significantly upregulated in diseased liver tissue, indicating their pivotal roles in processes such as apoptosis resistance, oxidative stress management, and metabolic reprogramming [51,52]. The increased expression of these markers correlates with enhanced tumor cell survival, angiogenesis, and tumorigenic progression in HCC. Conversely, downregulated genes reflect the reduced functional capacity of normal hepatocytes, further underscoring the metabolic and functional disruptions characteristic of liver cancer. Unexpectedly, certain markers like VCX2 exhibit lower expression levels in this dataset, which may lead to underestimation of their significance. This underscores the importance of analyzing expression data within broader statistical and biological contexts to ensure accurate marker interpretation [9].

The volcano plot in Figure 1D provides a broader view of differential gene expression, capturing the distribution of genes based on fold change and statistical significance. Highly upregulated genes, such as TMBIM4 and RGS5, are positioned on the right side of the plot, marking their pivotal roles in apoptosis inhibition, immune modulation, and angiogenesis—hallmarks of liver disease and cancer. Conversely, genes with significant downregulation on the left side likely represent those involved in hepatocyte-specific metabolic functions, which are suppressed during disease progression [53]. This visualization reinforces the critical roles of upregulated genes in diseased liver states, positioning them as potential biomarkers and therapeutic targets.

The expression patterns of the genes in both healthy and sick liver samples are shown in Figure 1E. The top three markers, RGS5, TMBIM4, and VCX2, are more prominent based on their percent expressed and average expression levels in the liver dataset. These markers are represented in the dot plot, where the size of the dots denotes the proportion of cells expressing each gene, and the color intensity reflects the average expression level. The expression patterns of these genes across the cell types are visible more clearly in the sick liver samples, where fibroblasts, macrophages, T cells, and B cells exhibit higher expression levels of TMBIM4, RGS5, and CEACAM7. TMBIM4 is particularly known for its function in apoptosis inhibition, supporting the survival of tumor cells and activated fibroblasts within the fibrotic and inflammatory environment of liver cancer [54]. This activity allows tumor cells to resist programmed cell death while thriving in the adverse conditions produced by the diseased liver.

HCC arises through a series of events, including chronic inflammation, immunological dysregulation, fibrosis, and extracellular matrix remodeling, all stimulating tumor proliferation [55,56]. Comprehending these pathways demands an extensive study of the cellular and molecular changes that trigger disease progression. Figure 2 illustrates a comprehensive and detailed analysis of the cellular and molecular differences between healthy and diseased liver tissue.

Figure 2A presents a bar plot illustrating the proportional distribution of liver cell types in healthy and diseased tissues, with distinct clusters representing different transcriptional fingerprints. In healthy liver, hepatocytes (green) dominate, reflecting their essential roles in metabolism, detoxification, and protein synthesis [57,58]. Minor populations of supporting cells, including macrophages (blue), endothelial cells (yellow), fibroblasts (brown), T cells (dark pink), and B cells (light pink), contribute to vascular integrity, immune surveillance, and extracellular matrix (ECM) regulation. In contrast, diseased liver samples exhibit a marked reduction in hepatocytes and an increase in fibroblasts, macrophages, and immune cells, indicating fibrosis, inflammation, and immune activation [59,60]. The expansion of these clusters highlights the progression from a functional liver architecture to a pathological state, with stromal and progenitor cells playing supportive roles in both conditions.

Building on these results, Figure 2B expands on these results by delineating enhanced signaling pathways in sick liver tissue, organized by gene ratio and statistical significance. The Phosphoinositide 3-Kinase – Protein Kinase B (PI3K-Akt) essential for cell survival, proliferation, and angiogenesis, is notably activated, facilitating tumor growth and apoptosis resistance [61,62]. Cytokine-cytokine receptor interactions are also enriched, reflecting the heightened inflammatory signaling driven by macrophages and T cells [63,64]. ECM-receptor interactions and focal adhesion pathways underline the role of fibroblasts and tumor cells in ECM remodeling and metastasis [65]. Additionally, ribosome-related pathways highlight the increased protein synthesis demands of proliferating tumor and stromal cells [66]. The activation of immune-related pathways, such as those associated with rheumatoid arthritis, indicates systemic immunological dysregulation in liver illness [67]. These findings highlight the interaction among tumor proliferation, fibrosis, immunological regulation, and angiogenesis in hepatic illness [68,69].

Figure 2C examines the biological processes increased in diseased liver tissue, providing functional insights into specific cell group activities. Chemotaxis the directed migration of immune cells like macrophages, T cells, and B cells to sites of inflammation, is significantly increased, reflecting chronic immune activation in liver disease and impacts the immune microenvironment in HCC [70]. ECM organization is driven by fibroblasts, which deposit collagen and other matrix components, perpetuating the fibrotic phenotype [71]. Dysregulated cell-substrate adhesion and increased cell adhesion indicate altered interactions between cells and the ECM, critical for tissue remodeling and tumor invasion. Heightened cytoplasmic translation and ribosomal activity further underscore the metabolic demands of tumor and stromal cells [72].

The cellular components implicated in liver pathology are highlighted in Figure 2D, illustrating structural and functional alterations during disease progression. Fibroblasts play a central role in extracellular matrix (ECM) remodeling, evidenced by the accumulation of collagen-rich matrices [73]. Enhanced focal adhesions and cell-substrate junctions facilitate tumor invasion and migration, while increased ribosomal activity and endoplasmic reticulum components reflect the elevated protein synthesis required for tumor progression [74,75]. Changes in the basement membrane, driven by endothelial and fibroblast cells, underscore the importance of these structures in tumor invasion and angiogenesis.

The molecular functions enriched in sick liver tissue are finally examined in Figure 2E, which examines the molecular processes enriched in pathological liver tissue, specifically with ECM-related activities. The increased production of extracellular matrix structural components, such as collagen, indicates the structural alterations caused by fibroblasts and activated hepatic stellate cells [76]. Integrin binding and cell adhesion molecule activity are crucial for cell motility, invasion, and tissue remodeling [77]. Elevated levels of heparin and glycosaminoglycan binding emphasize the modulation of growth factors and cytokine signaling, which drive angiogenesis and immune responses [78]. Additional processes, such as platelet-derived growth factor (PDGF) binding, reflect fibroblast activation and ECM remodeling [79].

The cellular and molecular reprogramming that happens during the progression of HCC and liver disease is comprehensively illustrated in this figure. The loss of hepatocytes and the growth of fibroblasts, macrophages, and immune cells represent significant architectural and functional changes in the liver microenvironment. Significant pathways include PI3K-Akt signaling, ECM-receptor interaction, and cytokine-cytokine receptor interaction demonstrating the relationship among tumor growth, fibrosis, immunological regulation, and angiogenesis. The biological processes of cell adhesion, extracellular matrix structure, and chemotaxis emphasize the active contributions of stromal and immune cells to the development of the fibrotic and inflammatory landscape of the diseased liver. Molecular mechanisms include platelet-derived growth factor (PDGF) signaling, ECM remodeling, and integrin binding highlighting the roles that stromal cells play in tumor growth and metastasis. These findings demonstrate the complexity of liver pathology and present important information on potential treatment targets for regulating tumor growth, immunological responses, and fibrosis in HCC and associated disorders.

The study extends its scope by analyzing the expression of liver cancer biomarkers across bladder, breast, colon, lung, ovarian, pancreatic, and gastric cancer. This comprehensive approach aims to assess whether these biomarkers are liver-specific or shared across other malignancies, providing critical insights into their diagnostic and therapeutic implications. By combining gene clustering, expression profiling, shared marker analysis, and statistical validation within the larger framework of tumorigenic processes, the study provides an in-depth picture of the transcriptional and functional landscape of liver cancer.

Figures 3A and 3B illustrate genes’ clustering and expression patterns across eight cancer datasets, emphasizing the molecular peculiarity of liver cancer and the common tumorigenic pathways among other malignancies. Figure 3A displays distinctive transcriptional fingerprints, with liver cancer constituting a separate cluster enriched in genes linked to apoptosis resistance, fibrosis, and vascular remodeling—characteristics of HCC progression [80]. Liver-specific clusters reflect the biological context of chronic liver injury, metabolic reprogramming, and inflammation, differentiating HCC from cancers like pancreatic, gastric, and pulmonary tumors. Shared pathways, such as ECM remodeling and immune modulation, are prevalent across cancers but lack organ-specific relevance [81].

**Figure 3.**
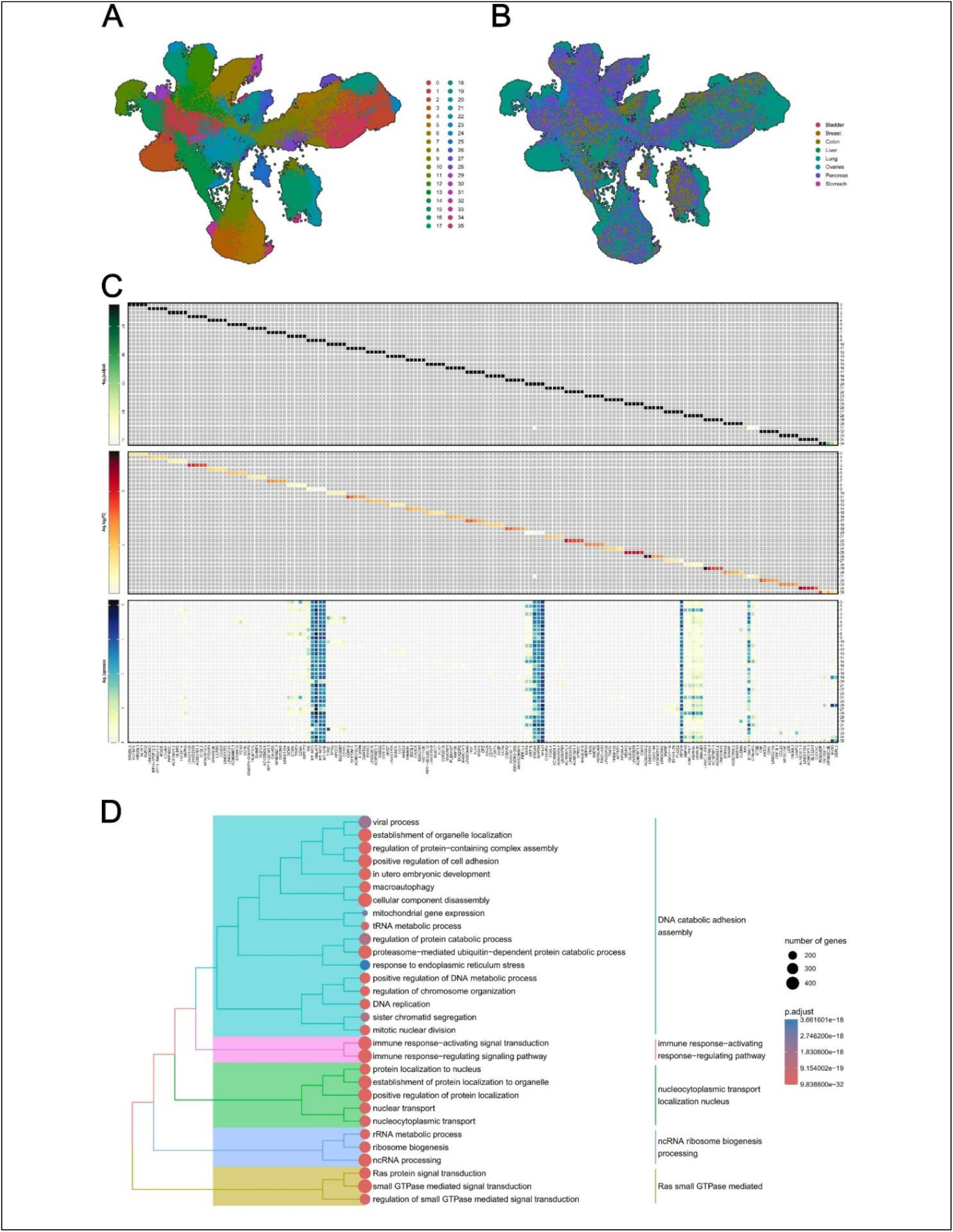
Comprehensive analysis of cancer biomarker expression and pathway enrichment across multiple cancer types. (A) and (B) present clustering maps derived from eight cancer datasets, identifying transcriptional patterns and distinct gene expression clusters associated with specific cancer types. (C) showcases a heatmap highlighting the top markers per cluster, illustrating their expression levels and functional roles across cancer datasets. (D) displays a tree plot of enrichment analysis for all biomarkers in the dataset, with pathways color-coded to represent their involvement in biological processes.

Figure 3B describes that liver-specific clusters exhibit markedly elevated expression levels in liver cancer relative to other malignancies, with genes regulating metabolism and fibrosis being minimally expressed in cancers such as breast or bladder, yet highly expressed in liver tissue. This reinforces the diagnostic capability of these clusters for liver cancer. This reinforces their diagnostic potential for liver cancer. Conversely, overlapping expression of genes involved in angiogenesis and ECM degradation across gastric and pancreatic tumors highlights shared tumorigenic pathways [82]. These findings highlight the simultaneous problems of distinguishing genuine liver-specific biomarkers while recognizing shared tumorigenic pathways.

To further investigate the specificity of liver cancer biomarkers across other malignancies, such as the bladder, breast, colon, lung, ovary, pancreas, and stomach cancer, Figure 3C presents a detailed heatmap analysis of gene expression across clusters derived from several cancer datasets. Key markers such as Metastasis-Associated Lung Adenocarcinoma Transcript 1 (MALAT1), Mitochondrially Encoded NADH Dehydrogenase 3 (MT-ND3), and Mitochondrially Encoded NADH Dehydrogenase 2 (MT-ND2), are prominently expressed across various cancers. MALAT1, a long non-coding RNA, is crucial for triggering metastasis, angiogenesis, and immunological modulation, significantly contributing to many malignancies [83]. Likewise, MT-ND3 and MT-ND2, involved in mitochondrial energy production, highlight metabolic adaptations common across tumors [84,85].

Markers include S100 Calcium-Binding Protein A6 (S100A6), S100 Calcium-Binding Protein A11 (S100A11), and Mitochondrially Encoded Cytochrome B (MT-CYB) also stand out for their roles in stress response, epithelial-to-mesenchymal transition (EMT), and metabolic reprogramming enhance tumor aggressiveness and survival. S100A6 and S100A11, constituents of the S100 protein family, are linked to cell migration and invasion, highlighting their role in metastasis [86,87]. MT-CYB, a mitochondrial gene associated with the electron transport chain, provides the metabolic requirements of rapidly multiplying tumor cells, confirming its significance in energy metabolism across many cancer types [88]. Additionally, ribosomal and metabolic genes like Glyceraldehyde-3-Phosphate Dehydrogenase (GAPDH), Ribosomal Protein L41 (RPL41), and Ribosomal Protein S29 (RPS29) reflect the biosynthetic and metabolic demands of tumor cells. GAPDH’s function in glycolysis, along with RPL41 and RPS29’s roles in protein synthesis, underscores their importance in supporting rapid tumor cell proliferation [89–91]. Interestingly, markers such as VCX2, while present in the dataset, exhibit lower expression compared to dominant markers like TMBIM4 and RGS5, which are highly expressed across multiple cancer types. TMBIM4’s role in apoptosis inhibition and RGS5’s involvement in vascular remodeling emphasize their significance in maintaining tumor microenvironments and supporting metastasis [11,92].

Figure 3D presents a tree plot that summarizes the enrichment analysis of biomarkers throughout the dataset, emphasizing pathways and processes linked to tumor growth. The identified pathways represent essential biological processes, including DNA catabolic adhesion, immune response activation and regulation, nucleocytoplasmic transport, nuclear localization, ncRNA processing, ribosome synthesis, and Ras small GTPase-mediated signaling. These processes show essential mechanisms that cause cancer, significantly influenced by markers such as TMBIM4, RGS5, and MALAT1, which are crucial in apoptotic regulation, vascular remodeling, and immunological modulation [93].

The figure is color-coded to represent each pathway’s statistical significance (adjusted p-value), with darker colors indicating greater significance. The size of the dot correlates to the number of genes involved in each process, emphasizing the extent to which these pathways are represented in the dataset. Pathways associated with DNA catabolism and nucleocytoplasmic transport exhibit a significant number of expressed genes, underscoring their extensive role in tumor cell proliferation and genomic instability [94]. Likewise, non-coding RNA (ncRNA) processing and ribosome biogenesis pathways are enhanced by markers such as RPL41 and RPS29, showing the increased requirement for protein synthesis in cancer cells [95–97].

With markers like CEACAM7 and MALAT1 enabling immune response-activating and regulating pathways, the plot also highlights the significance of immune regulation and signaling pathways [98]. These processes significantly influence tumor-immune interactions, immune evasion, and the development of a pro-inflammatory environment that promotes tumor growth. Furthermore, its function in cellular signaling, angiogenesis, and metastasis is reflected in the Ras small GTPase-mediated signaling pathway, which is triggered by markers like RGS5 [99].

Overall, Figure 3D provides a visually clear and statistically robust overview of the pathways enriched across cancer types, illustrating the number of genes expressed in each process and their relative significance. The tree plot not only highlights the shared tumorigenic mechanisms, such as immune modulation and ribosome biogenesis but also reveals the distinct molecular activities of specific cancer-related pathways, as demonstrated by the size and color coding of the enriched processes. This integrative analysis underscores the complex interplay of transcriptional networks driving cancer progression.

Building on this analysis, Figures 4A*–4G* present a detailed comparison of DEGs in liver cancer and seven other cancer types, highlighting the molecular distinctiveness of HCC. For instance, Figure 4A reveals an increased expression of Mitochondrial 2-Like 1 (MTRNR2L1), reflecting mitochondrial stress responses in HCC, and Apolipoprotein A2 (APOA2), a marker indicative of liver-specific lipid metabolism reprogramming [100,101]. In contrast, the significant downregulation of Albumin (ALB) in liver cancer, compared to bladder cancer, underscores the reduced synthetic capacity of diseased liver tissue [102]. An immune-related marker called Immunoglobulin Heavy Chain Protein (IGHGP) emphasizes how immunological responses contribute to the development of liver cancer [103]. Figure 4B compares breast and liver cancers, demonstrating higher expression of X-Inactive Specific Transcript (XIST), an epigenetic regulator, in breast cancer. Meanwhile, liver cancer exhibits an elevated expression of APOA2 and IGHGP, reinforcing its dependence on metabolic and immune regulatory pathways [104].

**Figure 4.**
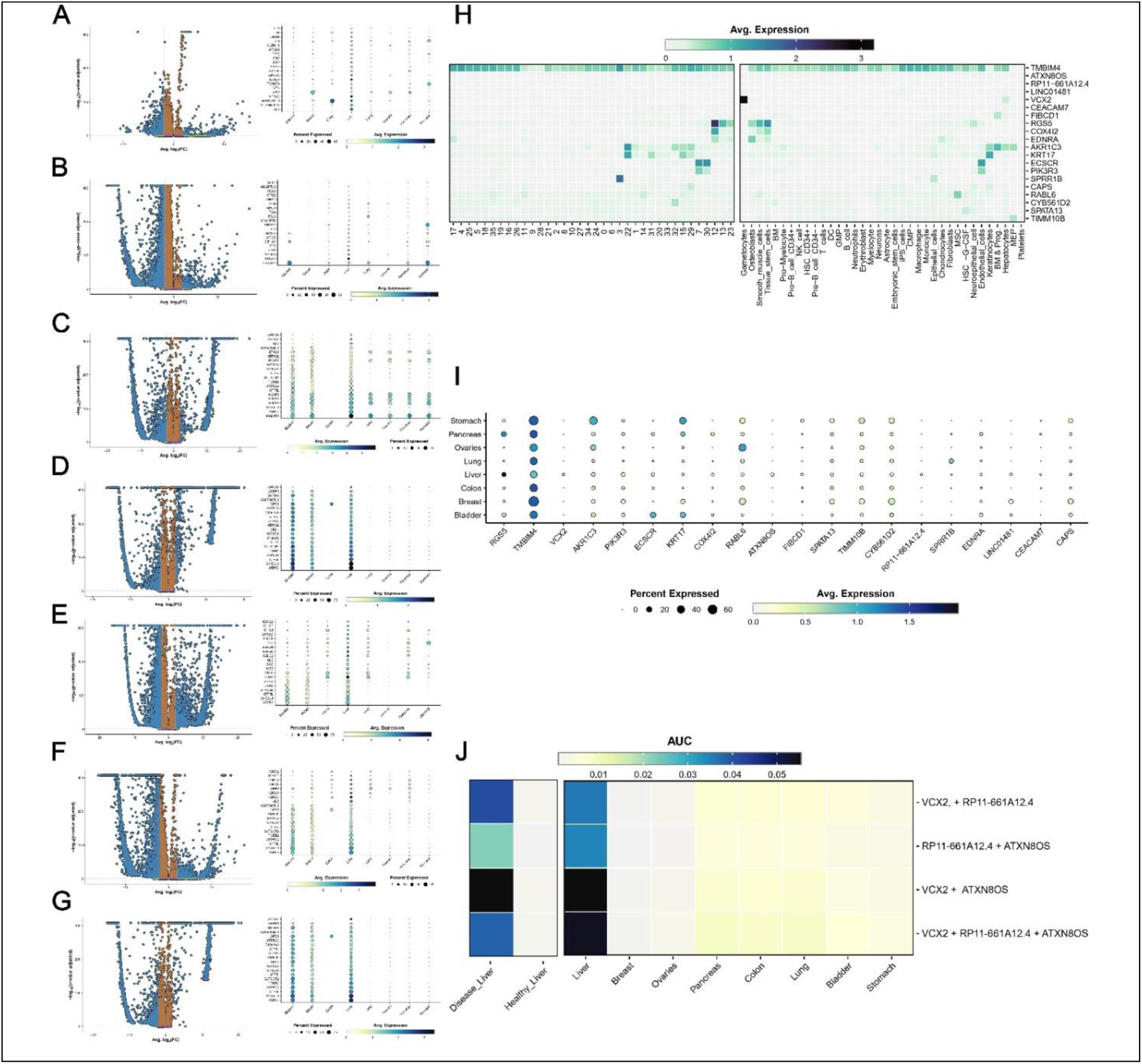
Comprehensive analysis of biomarkers and transcriptional changes in cancer tissues. The volcano plots show differentially expressed genes in liver cancer than (A) bladder, (B) breast, (C) colon, (D) lung, (E) ovaries, (F) pancreas, and (G) stomach cancers. The dot plots represent the results of the top 20 genes per comparison. (H) heatmap showing the results for each cluster and cell annotations for the genes expressed differently in HCC vs healthy liver, and (I) the dot plot of the same genes by cancer organ dataset. (J) enrichment heatmap for a combination of markers against healthy and diseased liver datasets and the eight cancer tissue datasets.

When liver and colon cancers are compared in Figure 4C, colon cancer exhibits higher levels of TFF1 and TFF2 (Trefoil Factors), which are indicators of epithelial maintenance and mucosal healing [105]. On the other hand, liver cancer shows increased expression of ALB (Albumin), APOA2 (Apolipoprotein A2), and IGHG3 (Immunoglobulin Heavy Constant Gamma 3), indicating liver-specific B-cell-mediated immune responses and systemic metabolic alterations [106]. Figure 4D compares liver and lung cancers, highlighting the elevated expression of Eukaryotic Translation Initiation Factor 1A, Y-linked (EIF1AY) in lung cancer, reflecting enhanced protein synthesis [107]. In liver cancer, the pronounced expression of fibrinogen components (FGA, FGB, FGG) and Transthyretin (TTR), a thyroid hormone transport protein, underscores its hypercoagulable state and metabolic dysregulation [108].

The role of key regulators in HCC progression becomes evident when examining angiogenesis and immune interactions. In the same way, RGS5 shows significant expression in endothelial cells and macrophages, contributing to angiogenesis and vascular remodeling [109]. These processes are vital for tumor progression, ensuring a continuous supply of oxygen and nutrients to rapidly growing cancerous tissue. The important role of RGS5 in the tumor microenvironment, especially in HCC, can be seen by its ability to mediate these processes. VCX2, while expressed at a lower ratio in certain cell types relative to TMBIM4 and RGS5, shows a unique expression pattern in the diseased liver. It contributes to chromosomal stability and may indirectly enhance tumor cell proliferation and survival [110]. Although its expression levels are slightly lower, its coexistence with the other two markers suggests a possible synergistic function in liver tumor biology. CEACAM7, widely expressed in immune cells, further supports this profile by increasing inflammation and facilitating immune evasion. Its function in enhancing inflammatory responses can produce a pro-tumorigenic environment, allowing cancer cells to evade immune detection [111].

The higher expression of GPX1 (glutathione peroxidase 1), which is essential for regulating oxidative stress in liver cancer [112], is highlighted in Figure 4E, which compares liver and ovarian cancer. Conversely, ovarian cancer shows a high expression of XIST, reflecting differences in epigenetic regulation between these malignancies. Liver cancer-specific markers, including APOA2 and IGHGP, further highlight its distinct immunological and metabolic landscape.

Figure 4F examines liver and pancreatic cancers, revealing elevated expression of ATP Synthase Subunit E (ATP5E) and Guanine Nucleotide-Binding Protein Subunit Beta-2-Like 1 (GNB2L1) in liver cancer, which support its energy demands through mitochondrial reprogramming [113]. While pancreatic cancer relies more on SPP1 (Osteopontin), a biomarker associated with ECM remodeling and metastasis, it is also moderately upregulated in liver cancer [114]. In Figure 4G, liver and stomach cancers are compared. Stomach cancer displays elevated TFF1 and TFF2, markers linked to mucosal defense, whereas liver cancer exhibits increased ALB, APOA2, and immune-related markers such as IGHG1 and fibrinogen subunits, reflecting its inflammatory and pro-coagulant tumor microenvironment.

Figure 4H showcases the expression levels of key markers across various cell types, where darker colors indicate higher expression and larger dot sizes represent the percentage of cells expressing each marker. TMBIM4 is widely expressed in nearly all cell types except megakaryocyte-erythroid progenitors, platelets, and gametocytes, highlighting its universal role in apoptosis inhibition and cellular stress regulation [92]. However, VCX2 is primarily found in gametocytes and hepatocytes, suggesting a specialized function in chromosomal stability and liver-specific processes [9]. RGS5 is prominently expressed in osteoblasts, smooth muscle cells, and tissue stem cells, and plays a key role in vascular remodeling, angiogenesis, and maintaining tumor microenvironments [49].

The presence of VCX2 in hepatocytes raises significant implications for its role in liver cancer biology. As a CTA, VCX2 is typically restricted to testicular germ cells, with its expression in somatic tissues being an aberrant event often driven by epigenetic dysregulation [9], epigenetic modifications contribute to immune evasion and tumor heterogeneity, which aligns with the potential role of VCX2 in HCC [115]. The study of Zhang et al. 2023 [116], further supports this, demonstrating that VCX2 expression is repressed in normal cells via DNA methylation but can be reactivated in cancer through DNA methyltransferase inhibitors (DNMTis) and histone deacetylase inhibitors (HDACis). Given this epigenetic plasticity, VCX2 might not only serve as a biomarker for HCC but also as a therapeutic target for epigenetic and immune-based interventions.

The cytoskeletal protein Keratin 17 (KRT17) mainly expressed in keratinocytes, supports epithelial integrity and tumor invasiveness, particularly in epithelial cancers [117]. Aldo-Keto Reductase Family 1 Member C3 (AKR1C3), involved in steroid metabolism, is highly expressed in the bone marrow and progenitor cells, reflecting its role in metabolic adaptation and differentiation [118]. Meanwhile, the Endothelial cell-specific chemotaxis receptor (ECSCR), a key endothelial marker, reinforces its importance in vascular signaling and homeostasis [119]. Though the heatmap highlights highly expressed markers, Ataxin 8 Opposite Strand (ATXN8OS) [120] and RP11-661A12.4 (lncRNA) [121], show lower expression, overshadowed by dominant markers like TMBIM4 and RGS5. Despite this, they may still play crucial roles in transcriptional regulation and specialized pathways, warranting further investigation.

In summary, Figure 4H highlights a diverse array of markers with distinct expression patterns across cell types. TMBIM4 is broadly expressed in nearly all cell types, emphasizing its universal relevance, while markers like VCX2, RGS5, KRT17, AKR1C3, and ECSCR [119] demonstrate more cell-type-specific expression, linked to specialized roles in gametocytes, hepatocytes, smooth muscle cells, keratinocytes, and endothelial cells, respectively. The emergence of VCX2 as a liver-specific marker in HCC further supports its relevance in oncogenesis, chromosomal stability, and tumor adaptation mechanisms, making it a compelling target for diagnostic, prognostic, and immunotherapeutic applications. This analysis also underscores the importance of exploring less prominently expressed markers, such as ATXN8OS and RP11-661A12.4, which may hold significant biological relevance despite their lower expression. The data highlights both the dominant transcriptional activity of key markers and the complexity of uncovering niche-specific pathways across diverse cell populations.

Figure 4I presents gene expression analysis across various cancer types and organs, revealing key statistical trends and marker distribution. Expression levels are assessed by average intensity (color gradients) and the percentage of cells expressing each marker (dot size). VCX2 shows strong, liver-specific expression, particularly in HCC, reinforcing its potential role in chromosomal regulation, immune evasion, and tumor progression. VCX2’s classification as a CTA makes it particularly attractive for immunotherapy, as its expression in tumors but not in most normal tissues minimizes off-target effects. The possibility of VCX2-directed therapies, including monoclonal antibodies, peptide vaccines, or even CAR-T cell therapy, aligns with strategies used for other CTA-based treatments.

Furthermore, TMBIM4 is broadly expressed across all organs and cancer types, reinforcing its role as a universal regulator of apoptosis and cellular stress response, essential for tumor survival [122]. In the liver, VCX2, RGS5, RP11-661A12.4, and ATXN8OS emerge as significant markers. VCX2 shows strong, liver-specific expression, particularly in HCC. RGS5, known for its role in vascular remodeling and angiogenesis, is also expressed in the liver, further supporting its involvement in tumor microenvironment modulation [123]. Additionally, RP11-661A12.4 and ATXN8OS, both lncRNAs, demonstrate liver-specific expression, suggesting potential regulatory roles in oncogenesis and positioning them as promising diagnostic biomarkers for HCC [124]. Their expression patterns highlight liver-specific carcinogenic mechanisms, including metabolic reprogramming, angiogenesis, and gene regulation, reinforcing their relevance in understanding liver cancer biology and identifying therapeutic targets.

Finally, Figure 4J examines the combinatorial expression of VCX2, ATXN8OS, and RP11-661A12.4 to identify synergistic interactions driving HCC progression. Analysis of marker pairings and the full triad in diseased liver tissue reveals that VCX2 + ATXN8OS exhibits the highest upregulation, suggesting that VCX2, a regulator of chromosomal stability and cell cycle progression, enhances ATXN8OS’s oncogenic role in transcriptional control. This reinforces the idea that VCX2 is not just a passive biomarker but a functional contributor to oncogenesis [125,126]. The VCX2 + RP11-661A12.4 combination also shows synergy, linking VCX2’s chromosomal regulatory function with RP11-661A12.4’s transcriptional activity, while ATXN8OS + RP11-661A12.4 displays functional overlap but a weaker effect. Interestingly, while the full triad (VCX2 + ATXN8OS + RP11-661A12.4) remains highly expressed, it does not surpass the intensity of VCX2 + ATXN8OS, emphasizing the distinct impact of this pairing on liver cancer biology. These findings highlight VCX2 as a potential master regulator in HCC development and a promising immunotherapeutic target. Future studies should focus on validating its role in tumorigenesis, further profiling its expression across cancer subtypes, and developing targeted therapies that leverage its unique tumor-associated expression.

Following the findings from Figure 4, Panel J, where VCX2 emerged as the most promising biomarker, further analysis confirmed its role in biological pathways and gene expression networks. Given its strong involvement in chromosomal stability, transcriptional control, and tumor progression, we extended our study by evaluating VCX2’s molecular interactions and its potential as a therapeutic target. The synthesis of these findings throughout Figure 4 highlights the intricate relationship between transcriptional reprogramming and cellular alterations in liver disease progression. The decrease in hepatocyte clusters and the proliferation of fibroblasts, macrophages, and immune cells indicate the pathological restructuring of the hepatic microenvironment, influenced by chronic damage, fibrosis, and carcinogenesis. Key genes such as TMBIM4 and RGS5 are identified as important regulators of these processes, positioning them as potential biomarkers and therapeutic targets. This study utilizes single-cell transcriptomics to elucidate the molecular mechanisms underlying liver pathology, facilitating precision medicine strategies aimed at common pathways in cancer progression.

To achieve this, we first examined VCX2’s relationship with other pathways and gene/protein networks, visualized in Figure 5, the upper part of the figure, which consists of three network diagrams. Figure 5A presents a node-link structure showing human liver expression data, where VCX2 (light pink) connects to multiple interacting genes or proteins, represented in shades of blue and gray. These interactions suggest functional relationships between liver cancer or chromosomal regulation networks, and different expressions of the network components in the liver tissue.

**Figure 5.**
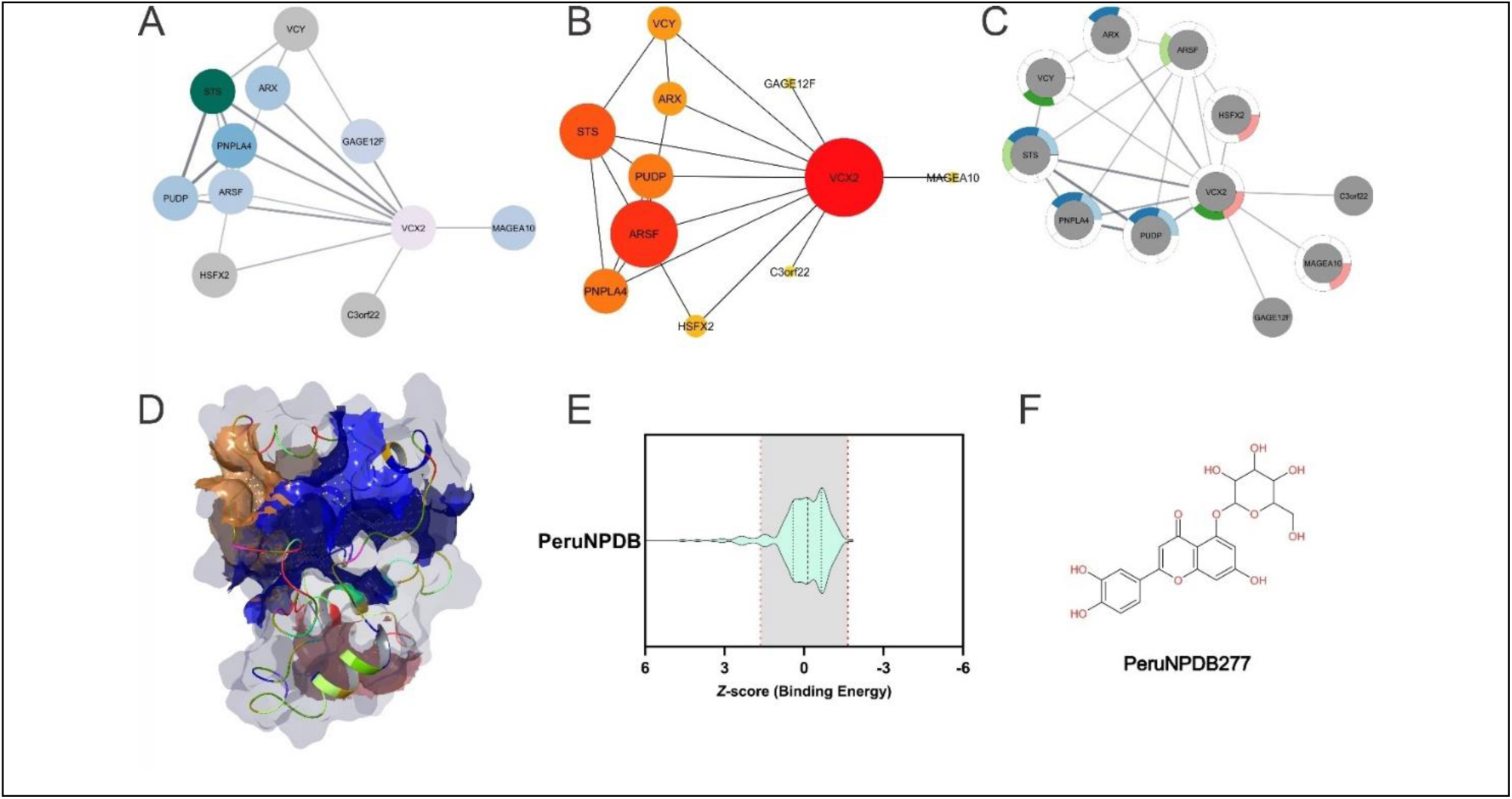
Network analysis and therapeutic exploration of VCX2. (Top) Three network diagrams illustrating VCX2’s molecular interactions. (A) Node-link diagram depicting VCX2 (pink) and its associated genes. (B) Centralized network emphasizing key partners. (C) Expanded interaction network, where color gradients highlight shared genes or proteins. (D) Identification of druggable pockets in VCX2. (E) Molecular docking results and binding affinity analysis (Z-score) to assess candidate compounds. (F) Structural representation of PeruNPDB277 (Luteolin-5-O-glucoside).

Figure 5B diagram positions VCX2 as a central red node, with the highest centrality scores on the network, directly connecting to key genes, including STS, ARSF, PUDP, and PNPLA4 (orange and yellow). These genes are involved in critical biological pathways: Steroid Sulfatase (STS) participates in steroid metabolism, particularly estrogen regulation, which may influence cancer progression [127]; Arylsulfatase F (ARSF) contributes to glycosaminoglycan metabolism, essential for extracellular matrix integrity [128]; Pseudouridine 5’-Phosphatase (PUDP) plays a role in RNA modification and stability, impacting gene regulation and tumorigenesis [129]; and Patatin-Like Phospholipase Domain Containing 4 (PNPLA4) is linked to lipid metabolism, a process highly relevant in liver cancer due to metabolic alterations in tumor growth [130]. The node sizes may indicate the strength or significance of these interactions, reinforcing VCX2’s relevance in multiple regulatory pathways. The interaction network in Figure 5C highlights the complexity of VCX2’s molecular associations, illustrating its involvement in diverse biological processes such as chromosomal regulation, transcriptional control, lipid metabolism, and tumor progression [9]. Nodes with color gradients or segmentations indicate shared proteins or genes across different datasets, while VCX2 shares annotations with genes such as the Heat shock transcription factor family, X-linked 2 (HSFX2), and the Melanoma-associated antigen 10 (MAGEA10) related to GAGE and XAGE proteins which have been implicated in human cancers [131] reinforcing VCX2’s central role in cancer biology.

The three-dimensional structural model of VCX2, shown in Figure 5D, reveals well-defined binding pockets (blue, orange, and pink), suggesting viable ligand interaction sites that position VCX2 as a promising target for drug discovery. Among these, the blue pocket ranked as the most favorable (site score 1.009, Maestro Sitemap), and was prioritized for MD and binding energy assessments. This structural model, likely derived from homology modeling or AlphaFold predictions, provides valuable insights into ligand binding. However, its accuracy remains limited due to the lack of an experimentally resolved crystal structure. While computational models offer a preliminary understanding of potential druggable sites, MD simulations are necessary to assess their flexibility and induced fit behavior. Furthermore, obtaining high-resolution structural data through X-ray crystallography or cryo-electron microscopy (cryo-EM) remains essential for refining binding site predictions and enhancing docking accuracy [132].

A virtual screening campaign was conducted using the PeruNPDB to identify potential inhibitors, focusing exclusively on the best-ranked pocket and applying a binding energy cut-off of −2 kcal/mol. The binding energy Z-score distribution in Figure 5E visualizes the results, where negative Z-scores indicate stronger ligand-protein interactions. The shaded confidence interval represents the distribution of binding affinities, while dashed red lines likely denote thresholds for statistically significant binders. The screening revealed multiple compounds with a strong affinity for VCX2, supporting the potential for novel inhibitor discovery. Among these, Luteolin-5-O-glucoside (PeruNPDB277) emerged as the top candidate, positioned within the high-affinity region of the Z-score distribution. The chemical structure of Luteolin-5-O-glucoside, depicted in Figure 5F, reveals its classification as a flavonoid glycoside, widely found in plants such as *Cirsium maackii* and *Equisetum arvense* [133,134].

Recognized for its anti-inflammatory, antioxidant, and anticancer properties, its aglycone form (luteolin) has been extensively studied for its ability to inhibit tumor cell proliferation, induce apoptosis, suppress metastasis, and interfere with angiogenesis [135]. Notably, Equisetum arvense has been traditionally used in Peruvian medicine for its diuretic, wound-healing, and anti-inflammatory properties, aligning with the biological activities attributed to luteolin-5-O-glucoside. Structurally, its multiple hydroxyl (-OH) groups facilitate strong molecular interactions, primarily through hydrogen bonding with key residues in VCX2’s active site, enhancing its potential as a therapeutic compound. The best-ranked pocket underwent additional restricted and limited docking to refine the binding predictions further to identify the most active region. Luteolin-5-O-glucoside demonstrated a docking score of −7.42 kcal/mol from the screened compounds, confirming a strong interaction with the binding pocket. However, docking scores alone do not fully capture the thermodynamic stability of the ligand-protein complex. MMGBSA calculations were employed to address this, providing a detailed breakdown of binding free energy contributions.

The MMGBSA analysis further refined the evaluation of ligand-protein interactions by integrating solvation effects and key energetic components, including electrostatics, van der Waals forces, and solvation-free energy. The calculated binding free energy (ΔG_bind) for Luteolin-5-O-glucoside was −40.13 kcal/mol, with a Coulomb interaction energy of −45.97 kcal/mol, indicating a highly stable and energetically favorable interaction. These strongly negative free energy values suggest spontaneous binding with high affinity, reinforcing Luteolin-5-O-glucoside as a strong candidate for VCX2 targeting.

Energy decomposition analysis revealed that electrostatic interactions serve as the primary stabilizing force within the binding pocket, with strong Coulomb energy underscoring the role of hydrogen bonding and charge complementarity between the ligand and key VCX2 residues. Van der Waals interactions further contribute to ligand accommodation, ensuring a well-fitted and tightly bound complex, while desolvation effects promote binding stability, as indicated by the favorable solvation energy components. Collectively, these factors reinforce the thermodynamic feasibility of ligand binding and suggest a high probability of successful inhibition.

Key active site residues involved in ligand binding were identified in Figure 6, through docking analysis, with Luteolin-5-O-glucoside forming strong hydrogen bonds with PRO91, GLU97, and GLU109 (Figure 6A), stabilizing its position within the VCX2 active site. Additionally, ARG46, ARG49, ARG50, LYS34, THR35, VAL38, PRO26, SER24, PRO94, GLU93, and GLU92 play significant roles in the ligand-protein interaction network, further contributing to binding stability (Figure 6B).

**Figure 6.**
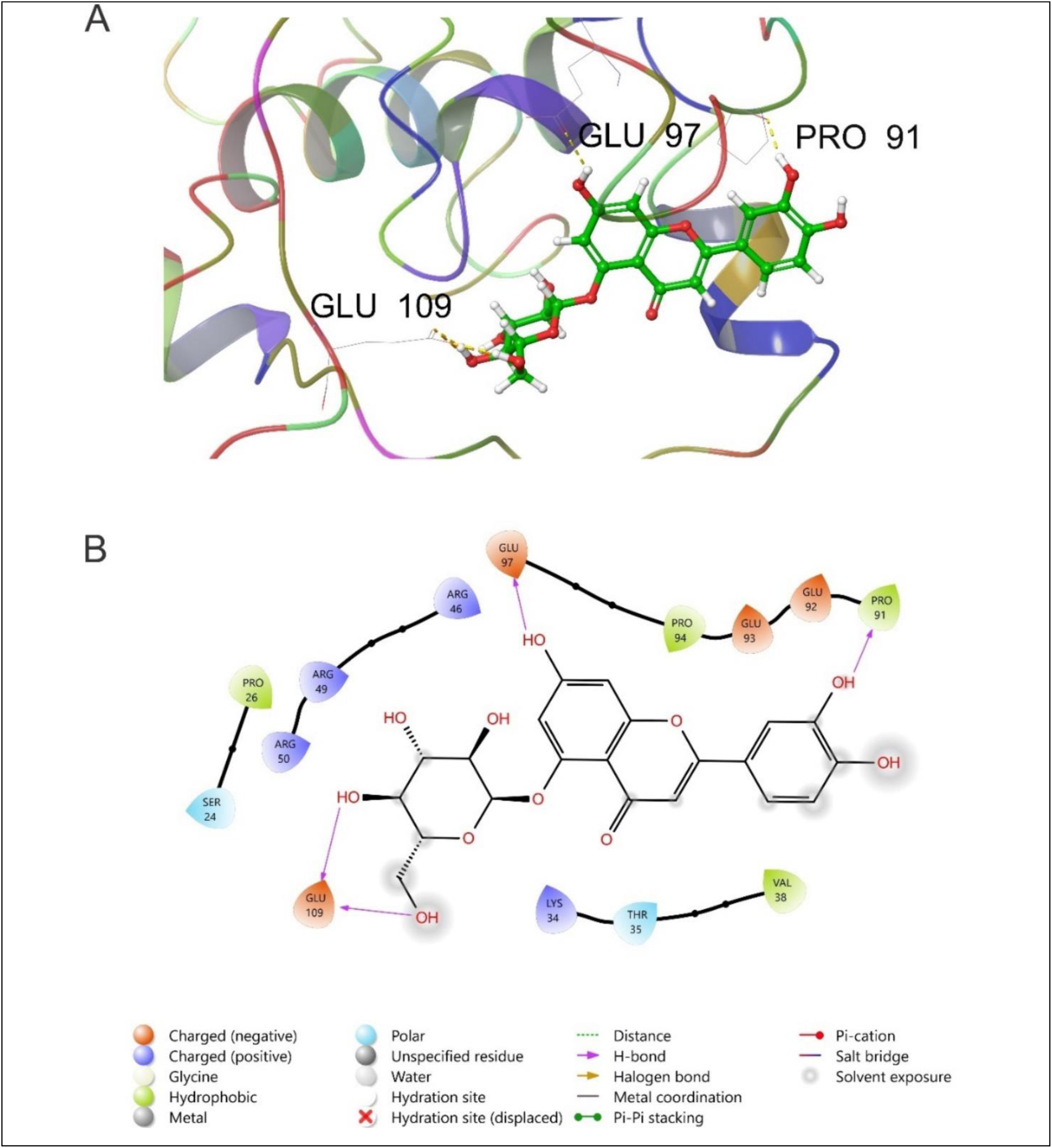
Molecular interactions of Luteolin-5-O-glucoside with VCX2. (A) 3D binding conformation of Luteolin-5-O-glucoside (green) within the VCX2 active site, showing hydrogen bonds (yellow dashed lines) with PRO91, GLU97, and GLU109. (B) 2D interaction map highlighting key ligand-protein interactions, including hydrogen bonding, pi-pi stacking, and electrostatic interactions.

To explore the dynamic behavior of Luteolin-5-O-glucoside within the VCX2 binding pocket, MD simulations were planned. MD simulations provide critical insights into ligand stability, conformational flexibility, and long-term interaction patterns. However, the absence of an experimentally resolved crystal structure posed significant challenges. The VCX gene family, including VCX2, has been implicated in certain cancers, although its precise role remains under investigation. While VCX genes are primarily expressed in male germ cells, emerging studies suggest their aberrant expression in cancer cells could serve as a biomarker or therapeutic target [136]. Key residues identified in this study, such as GLU97, PRO91, and GLU109, are conserved across related VCX family proteins, suggesting their role in ligand interactions and oncogenic activity. The identification of Luteolin-5-O-glucoside as a VCX2-binding compound suggests that it may disrupt VCX2-associated pathways, potentially interfering with oncogenic signaling mechanisms. The strong binding affinity, favorable free energy calculations, and interaction with key VCX2 residues reinforce its potential as a novel small-molecule inhibitor for VCX2-expressing cancers. Future studies should focus on *in vitro* and *in vivo* validation, SAR studies, and optimization of lead compounds for improved efficacy and drug-like properties.

## 4. Conclusion

This study was designed to identify a specific HCC biomarker, leading to the selection of VCX2 as a promising candidate. Through scRNA-seq analysis and transcriptomic profiling, we determined that VCX2 exhibits differential expression in HCC, distinguishing healthy liver tissue from diseased states. To assess the uniqueness of VCX2 as an HCC-specific marker, we expanded our analysis across multiple cancers, evaluating its expression in bladder, breast, colon, lung, ovarian, pancreatic, and stomach cancers. These findings validated VCX2 as a potentially unique biomarker for HCC, with implications for its role in tumor progression, chromosomal regulation, and oncogenic adaptation.

Following this multi-cancer validation, we performed an advance *in silico* studies to explore VCX2’s potential as a therapeutic target. Using structural modeling, MD, and virtual screening, we searched for natural products that could modulate VCX2 activity. The FASTA sequence of VCX2 was retrieved from AlphaFold and modeled, and its druggability was assessed using Maestro Sitemap, which identified multiple potential binding pockets. The most favorable binding site (blue pocket, site score 1.009) was selected for high-precision docking against the PeruNPDB natural product database.

Among the screened compounds, Luteolin-5-O-glucoside (PeruNPDB277), from *Equisetum arvense*, emerged as the top candidate, demonstrating a strong binding affinity with a docking score of −7.42 kcal/mol. To refine our predictions, Molecular Mechanics Generalized Born Surface Area (MMGBSA) analysis was employed, calculating a binding free energy (ΔG_bind) of −40.13 kcal/mol and a Coulomb interaction energy of −45.97 kcal/mol, indicating a highly stable and spontaneous interaction. Key stabilizing interactions included hydrogen bonding with PRO91, GLU97, and GLU109, further reinforced by electrostatic and van der Waals contributions.

Given that VCX2 may act as an oncogenic driver, its inhibition via small molecules could provide a potential strategy to desensitize tumor cells, particularly if VCX2 plays a role in cancer progression or therapy resistance. However, due to the lack of an experimentally resolved crystal structure, structural models relied on homology modeling and AlphaFold predictions, which, while informative, require further refinement. To enhance the reliability of these computational predictions, MD simulations were planned but faced with limitations due to structural uncertainties. Future studies should aim for experimental structural characterization (X-ray crystallography or cryo-EM) and in vitro validation of VCX2-targeting compounds to confirm their therapeutic potential.

In conclusion, this study provides a comprehensive pipeline integrating single-cell transcriptomics, multi-cancer validation, high-precision molecular modeling, and virtual screening to explore VCX2 as a novel HCC biomarker and drug target. The identification of Luteolin-5-O-glucoside as a lead compound highlights its potential for targeted therapy, opening avenues for precision medicine approaches aimed at modulating VCX2’s role in tumor progression. Future research should focus on experimental validation and structure-based drug design to optimize inhibitors that can modulate VCX2-driven oncogenic pathways in HCC and potentially other cancers.

## Funding

This work was funded by the Consejo Nacional de Ciencia, Tecnología e Innovación Tecnológica (CONCYTEC), the Programa Nacional de Investigación Científica y Estudios Avanzados (PROCIENCIA), by the call “E067-2023-01 Proyectos Especiales: Proyectos de Incorporación de Investigadores Postdoctorales en Instituciones Peruanas” [número de contrato PE501084367-2023], and Grant PE501088204-2024. Also, it was funded by the Universidad Católica de Santa Maria (grants 27499-R-2020, 27574-R-2020, 7309-CU-2020, and 28048-R-2021) and by the Research Management Office from the Universidad Católica de Santa María.

## Institutional Review Board Statement

Not applicable.

## Informed Consent Statement

Not applicable.

## Data Availability Statement

The software used in this study includes Maestro (Schrödinger Suite), Desmond, SiteMap, and Glide, all of which require a paid license for both academic and commercial use. Additionally, STRING, Cytoscape, and PASSer were utilized, which are freely available for non-commercial use. All software tools are referenced throughout the Methods and Results sections.

Furthermore, protein structures, docking grids, molecular dynamics trajectories, ligand preparation files, energy minimization data, and analysis scripts generated during this study are documented accordingly. All simulation scripts, docking results, molecular interaction analyses, and processed datasets will be made available upon request or through a publicly accessible repository.

## Acknowledgments

The authors express their gratitude for the financial support from the Programa Nacional de Investigación Científica y Estudios Avanzados—PROCIENCIA (PE501088204-2024 and PE501084367-2023).

## Conflicts of Interest

The authors declare no conflicts of interest.

## List of Abbreviations

ALB: Albumin
AKR1C3: Aldo-Keto Reductase Family 1 Member C3
AFP: alpha-fetoprotein
APOA2: Apolipoprotein A2
ARSF: Arylsulfatase F
ATP5E: ATP Synthase Subunit E
ATXN8OS: Ataxin 8 Opposite Strand
BAX: BCL2 associated X, apoptosis regulator
BP: Biological Processes
CTA: cancer/testis antigen
CEACAM7: CEA Cell Adhesion Molecule 7
CC: Cellular Components
cryo-EM: cryo-electron microscopy
CYB5R3: Cytochrome B5 Reductase 3
DEG: differential gene expression
DEGs: Differentially expressed genes
DNMTis: DNA methyltransferase inhibitors
ECSCR: Endothelial cell-specific chemotaxis receptor
EMT: epithelial-to-mesenchymal transition
EIF1AY: Eukaryotic Translation Initiation Factor 1A, Y-linked
XP: Extra Precision
ECM: extracellular matrix
GEO: Gene Expression Omnibus
GO: Gene Ontology
GAPDH: Glyceraldehyde-3-Phosphate Dehydrogenase
GNB2L1: Guanine Nucleotide-Binding Protein Subunit Beta-2-Like 1
HSFX2: Heat shock transcription factor family, X-linked 2
HBV: hepatitis B virus
HCV: hepatitis C virus
HCC: Hepatocellular carcinoma
HTVS: High-Throughput Virtual Screening
HDACis: histone deacetylase inhibitors
IGHGP: Immunoglobulin Heavy Chain Protein
KRT17: Keratin 17
KEGG: Kyoto Encyclopedia of Genes and Genomes
MCC: Maximal Clique Centrality
MAGEA10: Melanoma-associated antigen 10
MALAT1: Metastasis-Associated Lung Adenocarcinoma Transcript 1
MTRNR2L1: Mitochondrial 2-Like 1
MT-CYB: Mitochondrially Encoded Cytochrome B
MT-ND2: Mitochondrially Encoded NADH Dehydrogenase 2
MT-ND3: Mitochondrially Encoded NADH Dehydrogenase 3
MD: molecular docking
MF: Molecular Functions
NAFLD: non-alcoholic fatty liver disease
ncRNA: non-coding RNA
PNPLA4: Patatin-Like Phospholipase Domain Containing 4
PeruNPDB: Peruvian Natural Products Database
PI3K-Akt: Phosphoinositide 3-Kinase - Protein Kinase B
PDGF: platelet-derived growth factor
PCA: principal component analysis
PPI: protein-protein interaction
PUDP: Pseudouridine 5’-Phosphatase
RGS5: Regulator of G-protein signaling 5
RPL41: Ribosomal Protein L41
RPS29: Ribosomal Protein S29
S100A11: S100 Calcium-Binding Protein A11
S100A6: S100 Calcium-Binding Protein A6
STRING: Search Tool for the Retrieval of Interacting Genes/Proteins
SMILES: simplified molecular-input line-entry system
scRNA-seq: single-cell RNA sequencing
SP: Standard Precision
STS: Steroid Sulfatase
SAR: structure activity relationship
TCM: Traditional Chinese Medicine
TMBIM4: Transmembrane BAX Inhibitor Motif Containing 4
TTR: Transthyretin
TM5B4X: Tropomyosin Beta Chain 4X
UMAP: Uniform Manifold Approximation and Projection
VCX2: Variable Charge X-Linked 2
XIST: X-Inactive Specific Transcript

## Notes

### Competing Interest Statement

The authors have declared no competing interest.

